# A Dynamic Framework for Context-Dependent Microbial Assembly: Validation of the Domain-Specific Stochastic–Deterministic Integration Model

**DOI:** 10.1101/2025.05.04.652157

**Authors:** Min-Tao Wan, Ping-Ju Liu

## Abstract

Microbial community assembly is governed by the interplay of stochastic and deterministic processes. However, most existing frameworks treat microbial communities as taxonomically homogeneous entities, overlooking the phylogenetic divergence and ecological distinctiveness of major microbial domains. Here, we present a new conceptual model—Domain-Specific Stochastic–Deterministic Integration (DSSDI)—that explicitly incorporates domain-level asymmetries, cross-domain interactions, and functional buffering into the community assembly process. Using 16S rRNA sequencing data from seasonal intertidal microbial communities, we tested four hypotheses derived from the DSSDI framework. Our results revealed that bacterial and archaeal communities differ in their stochastic–deterministic assembly patterns, spatial–temporal responses, network modularity, and functional robustness. Cross-domain cooperation emerged as a stabilizing factor, promoting metabolic complementarity and buffering against taxonomic turnover. This study introduces and validates the DSSDI framework as the first domain-explicit model of microbial community assembly, establishing a clear conceptual foundation and claiming priority for future theoretical and empirical developments in this field.

## Introduction

Microbial communities underpin critical global ecosystem functions—including carbon fixation, nitrogen transformation, and sulfur cycling—and are central to Earth’s biogeochemical cycles [1–3]. Despite their ecological importance, the mechanisms that govern microbial community assembly and the maintenance of functional resilience under environmental fluctuations remain incompletely understood.

Classical ecological theories, such as neutral and niche theory, have long provided foundational perspectives on community assembly. Neutral theory emphasizes stochastic processes like dispersal and ecological drift [4, 5], while niche theory focuses on deterministic forces such as environmental filtering and trait-based competition [6, 7]. Although influential, these frameworks often treat microbial communities as taxonomically and functionally uniform, thereby overlooking potential domain-specific assembly dynamics and interdomain interactions.

The stochastic–deterministic continuum model proposed by Zhou and Ning [8] was a conceptual advancement that integrated both forces across spatiotemporal scales. However, this model still assumes taxonomic and functional uniformity, overlooking the ecological divergence among microbial domains. Emerging evidence reveals that bacteria and archaea differ significantly in life-history strategies, metabolic flexibility, dispersal potential, and environmental tolerance [9, 10], suggesting that distinct assembly mechanisms may predominate within each domain.

Moreover, while the continuum framework advanced our understanding of community structuring, it largely neglects positive interdomain interactions—such as syntrophy, mutualistic exchange, and metabolic complementarity—that can enhance community stability and ecosystem resilience [11, 12]. Although Zhou and Ning acknowledged the role of species interactions, such dynamics were not formally integrated into their model. In the absence of interaction-explicit frameworks and domain-resolved analyses, existing models remain limited in explaining the complexity and stability of microbial ecosystems.

To fill this theoretical gap, we present the Domain-Specific Stochastic–Deterministic Integration (DSSDI) framework—a novel model that systematically integrates domain-level assembly dynamics with cross-domain interactions. DSSDI introduces a dual-axis structure: (i) domain-specific variation in the dominance of stochastic and deterministic processes, and (ii) the contribution of cross-domain cooperation to community modularity and functional buffering. The first axis posits that microbial domains (e.g., Bacteria and Archaea) exhibit differential sensitivity to ecological drivers due to their distinct evolutionary histories and physiological strategies. DSSDI rejects fixed assembly rules and instead asserts that process dominance is context-dependent, shaped by environmental heterogeneity, resource dynamics, and the presence or absence of interaction partners [8–10, 13]. The second axis highlights the role of positive interdomain interactions in fostering modular network structures, which in turn enhance system-level stability under environmental perturbation [11, 12, 14].

The DSSDI framework is grounded in four conceptual pillars (Figure 1):

(1) microbial community assembly is domain-specific and context-dependent;
(2) spatial and environmental heterogeneity modulate assembly strength across ecological scales [7, 15–16];
(3) cross-domain cooperation actively promotes network modularity and stability; and
(4) functional stability emerges not solely from taxonomic redundancy, but through taxonomic–functional decoupling and interdomain complementarity [17–19].

**Fig. 1.**
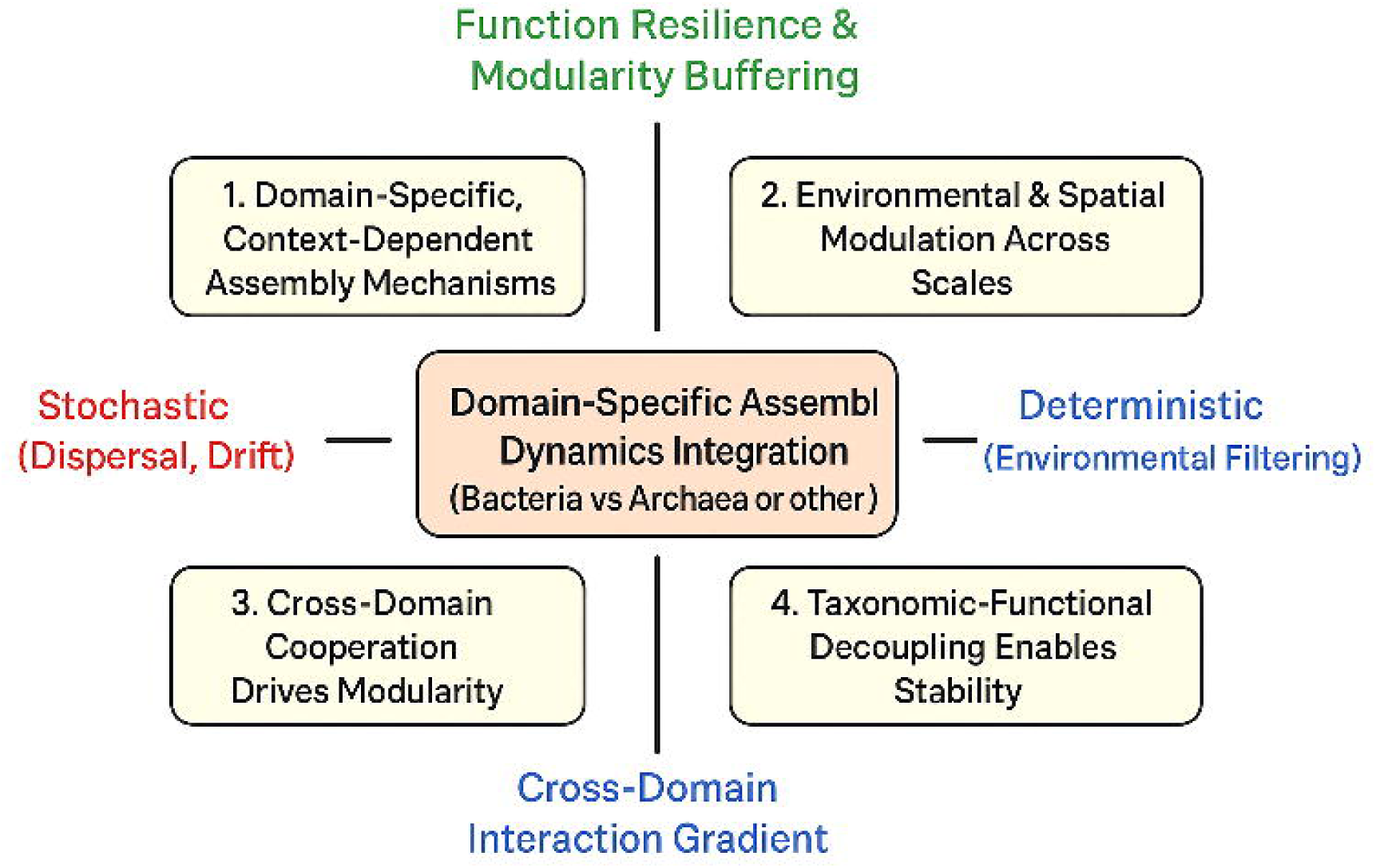
Conceptual framework of Domain-Specific Stochastic–Deterministic Integration (DSSDI). This framework illustrates the dual influence of stochastic processes (e.g., dispersal and ecological drift) and deterministic processes (e.g., environmental filtering) in shaping microbial community assembly, particularly for bacteria and archaea. The central box represents domain-specific assembly dynamics, which vary depending on phylogenetic domain and ecological context. The vertical axis captures the gradient toward functional resilience and modularity buffering, while the horizontal axis reflects the cross-domain interaction gradient. The four surrounding components denote the major hypotheses: (Top-left) domain-specific, context-dependent mechanisms govern microbial structuring; (Top-right) environmental and spatial drivers operate across multiple ecological scales; (Bottom-left) cross-domain cooperation enhances community modularity; and (Bottom-right) taxonomic– functional decoupling contributes to overall ecosystem stability.

To our knowledge, this is the first theoretical framework that explicitly incorporates domain-level assembly asymmetry and cross-domain ecological interactions into a unified model. By formally articulating and empirically validating the DSSDI framework, this study establishes conceptual and empirical priority in the development of a domain-explicit theory of microbial community assembly.

We evaluated DSSDI using microbial data from a dynamic intertidal aquatic system influenced by tidal, seasonal, and riverine gradients—conditions that jointly shape dispersal, environmental filtering, and interdomain cooperation. Within this setting, we tested four linked hypotheses:

(H1) Bacterial and archaeal communities exhibit domain-specific and context-dependent assembly, with shifts in the relative strength of stochastic versus deterministic processes;
(H2) Spatial, temporal, and environmental gradients jointly determine microbial composition at fine scales;
(H3) Cross-domain cooperation contributes to modular network structures that buffer taxonomic turnover; and
(H4) Functional complementarity across domains supports system-level function through taxonomic–functional decoupling.

By integrating classical ecological theory with domain-resolved ecological mechanisms and cooperative network principles, DSSDI provides a scalable, mechanistically grounded, and empirically testable model to explain how microbial communities assemble, interact, and persist across fluctuating environments. It offers a robust conceptual foundation for unifying microbial ecology across natural and engineered systems.

## Materials and Methods

To enhance readability, a condensed overview of methods is provided here. Full methodological details are available in Supplementary Methods (Supplementary Information, Section S1).

### Study Area and Sampling Strategy

Sampling was performed at nine fixed intertidal sites along the Nanmen coastline, Kinmen Island (Taiwan Strait), from July to November 2022, capturing seasonal variability driven by tides, monsoons, and riverine input. At each monthly time point (five total), a composite water sample (∼2 L) was collected per site by pooling six 1-meter interval subsamples along a 6 m linear transect centered at each GPS-marked location (45 total samples). This design minimized microscale heterogeneity and ensured representative sampling.

### Environmental Measurements

In situ water parameters—including temperature, pH, salinity, dissolved oxygen (DO), oxidation–reduction potential (ORP), total dissolved solids (TDS), and conductivity—were recorded using multiparameter probes (U-50, Horiba; VZ8403BZ, Hengxin). These variables were used in multivariate analyses, including db-RDA and variation partitioning, to assess environmental contributions to microbial community variation. Nutrient concentrations such as nitrate, phosphate, or ammonium were not measured in this study. We acknowledge this limitation and recommend that future studies incorporate nutrient data to improve the resolution of environmental filtering models and disentangle potential biogeochemical drivers of microbial assembly.

### DNA Extraction and Amplicon Sequencing

Environmental DNA was extracted from 0.22 μm PES filters using the DNeasy PowerWater Kit (Qiagen). Separate 16S amplicons were generated for bacteria (V6– V8, 968F–1391R) [20] and archaea (V3–V4, A306F–A712R) [21], using Phusion High-Fidelity polymerase. Barcoded amplicons were quantified (Qubit 2.0) and pooled equimolarly for sequencing.

### Sequence Processing and Normalization

Paired-end reads were merged (USEARCH v11.0) [22] and denoised into ASVs using UNOISE3 [23]. Chimeras were removed, and exact-sequence ASVs retained. To address domain-specific sequencing depth and preserve within-domain diversity, samples were rarefied separately for bacteria (2,352 reads/sample) and archaea (21,735 reads/sample). All downstream analyses—including diversity metrics, community composition, and assembly modeling—were performed independently within each domain and internally standardized. Cross-domain comparisons (e.g., interaction networks, functional stability) were based on topological or correlation-based metrics, rather than direct diversity indices, thereby minimizing bias introduced by differential rarefaction.

### Taxonomic Classification and Functional Prediction

ASVs were assigned taxonomy using SINTAX (SILVA 138 NR99, confidence ≥0.8) [24, 25]. Functional profiles were inferred with PICRUSt2 (KEGG Orthologs) [26], and centered-log ratio (CLR) transformed prior to ordination. Archaeal predictions were interpreted cautiously due to reference limitations.

### Diversity and Statistical Analyses

Alpha diversity (Shannon, richness) was computed from rarefied tables [27]. One-way ANOVA or Kruskal–Wallis tests assessed spatiotemporal differences. Beta diversity was assessed via Bray–Curtis dissimilarities and NMDS ordination [27]; PERMANOVA [28] and betadisper [29] tested group differences (9,999 permutations).

### Community Assembly and Environmental Structuring

Sloan’s neutral model [5] was fitted per domain to estimate stochasticity (R², immigration rate m), with bootstrapped confidence intervals. db-RDA [30], variation partitioning [31], and local contributions to beta diversity (LCBD) [32] analyses (Hellinger-transformed) assessed environmental and spatial effects [27] and adespatial packagesp [33].

### Network Construction and Core Taxa Identification

To ensure comparability across domains, we selected an equal number of taxa from each domain as network nodes. Specifically, all 39 archaeal genera identified in the dataset were included, and the top 39 most abundant bacterial genera (based on average relative abundance across all samples) were chosen to match. These bacterial taxa alone accounted for 76.7% to 93.3% of the total reads in each sample (pre-rarefaction), indicating strong ecological representativeness and community dominance. This approach minimized noise from rare taxa while preserving the core ecological structure for robust network inference. Genus-level co-occurrence networks were inferred using SparCC [34], SPIEC-EASI [35], and FlashWeave [36]. Consensus edges (≥2 methods) were retained. Networks were analyzed for degree, k-core, and Louvain modularity [37–39]. Module robustness was evaluated via bootstrap resampling (n = 20). Core taxa were defined by consistent high centrality.

### Functional Stability Simulation

To simulate disturbance, the lowest 30% of mean-abundance ASVs were removed per sample (100 iterations), and Bray–Curtis distances compared functional profiles before and after perturbation. Functional co-occurrence networks were also constructed to detect cross-domain metabolic modules [26].

### Software and Visualization

All analyses were performed in R v4.3.1 using packages including vegan [27], adespatial [33], phyloseq [40], igraph [37], picante [41], ggplot2 [42], and ggpubr [43]. Statistical significance was set at *p* < 0.05.

## Results

### Sequencing Data Summary

High-throughput amplicon sequencing of bacterial and archaeal communities generated a total of 9,494,093 raw paired-end reads. Among these, 5,592,468 reads originated from bacterial libraries, and 3,901,625 from archaeal libraries. After merging paired-end reads, 68.34% of bacterial and 78.11% of archaeal sequences were retained, yielding 3,410,000 high-quality bacterial and 2,860,000 high-quality archaeal reads following quality control. Dereplication and denoising produced 5,350 bacterial ASVs and 3,870 archaeal ASVs.

### Overview of Microbial Community Structure

To establish the baseline understanding of microbial community dynamics within the studied intertidal ecosystem, we characterized the taxonomic composition, spatiotemporal variability, and alpha diversity patterns across bacteria and archaea. At the phylum level, bacterial communities were dominated by Proteobacteria (74.2% ± 4.9%), followed by Cyanobacteria (10.8% ± 3.3%) and Bacteroidetes (3.6% ± 0.5%) (Fig. 2A), whereas archaeal assemblages were overwhelmingly composed of Thaumarchaeota (66.2% ± 7.4%) and Euryarchaeota (32.8% ± 7.9%) (Fig. 2B). This taxonomic profile suggests a higher compositional complexity within bacterial communities, contrasted by a narrower archaeal community structure centered around a few dominant lineages.

**Fig. 2.**
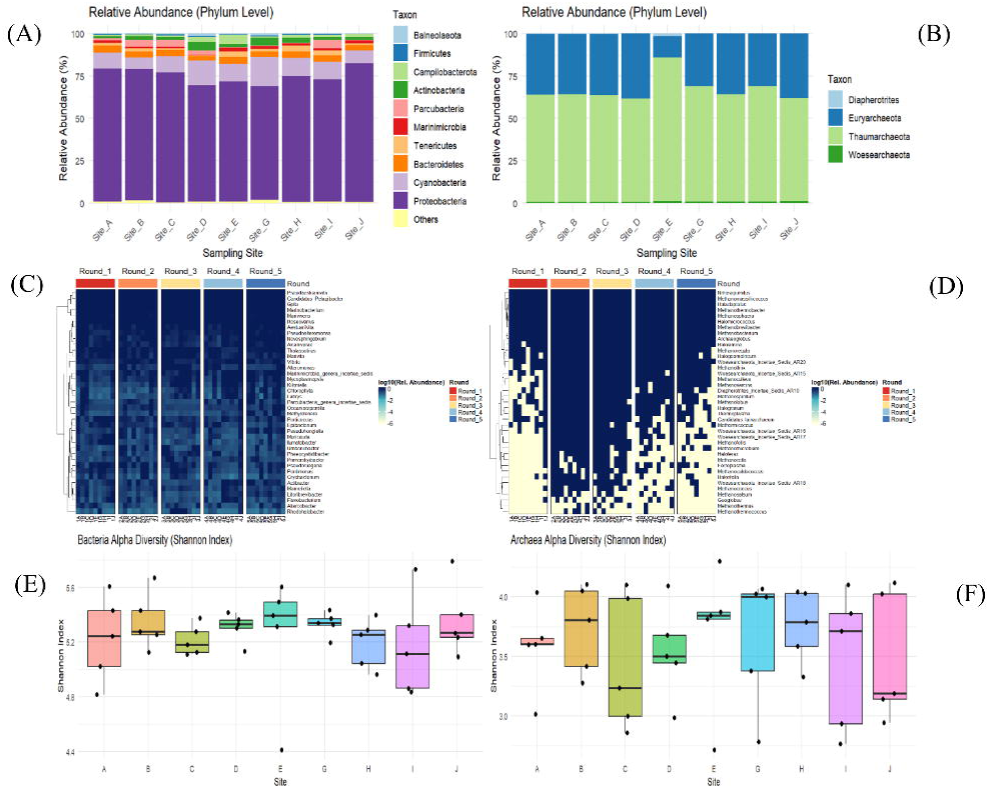
Taxonomic composition and alpha diversity of bacterial and archaeal communities. (A–B) Phylum-level relative abundances of bacterial (A) and archaeal (B) communities. (C) Genus-level relative abundances of dominant bacterial taxa across sampling times and locations. (D) Genus-level relative abundances of archaeal taxa across samples. (E–F) Shannon diversity indices for bacterial (E) and archaeal (F) communities across sampling sites.

Spatial and temporal patterns further highlighted contrasting dynamics between domains. Bacterial genera exhibited pronounced spatial heterogeneity and temporal turnover, with site-specific shifts (e.g., GpIIa surged to 34.5% relative abundance at Site G in October) and seasonal transitions (Pseudaestuariivita dominated early months, declining after August) (Fig. 2C). In contrast, archaeal communities were consistently dominated by Nitrosopumilus (62.2% ± 8.9%) across sites (Fig. 2D), with synchronized temporal fluctuations — peaking between July and September and declining toward November — suggesting tighter environmental regulation.

Alpha diversity analyses indicated that both bacterial and archaeal communities maintained relatively stable within-sample diversity across spatial gradients. Bacterial Shannon indices ranged from 4.57 to 5.59 (Fig. 2E), while archaeal Shannon indices ranged from 3.31 to 4.60 (Fig. 2F). One-way ANOVA and Kruskal-Wallis tests revealed no significant differences in alpha diversity across sites for either domain (p > 0.05 for all tests), despite observed spatial and temporal environmental variability.

Together, these findings show that bacterial communities exhibit higher spatial and temporal variability, while archaeal communities display greater compositional dominance and seasonal consistency. These contrasting patterns validate the hypothesis that domain-specific assembly processes shape microbial dynamics and enable testing DSSDI predictions on stochastic–deterministic balance and functional modularity.

### Domain-Specific Assembly: Stochastic and Deterministic Processes

To evaluate whether microbial communities conform to the predictions of neutral theory, we performed a three-tiered analytical approach. At the descriptive level, we constructed rank-abundance plots (RAPs) based on total ASV abundances across all samples for both bacterial and archaeal communities (Fig. 3A). Both domains exhibited classic long-tailed distributions, consistent with the neutral theory expectation of “a few dominant taxa and many rare ones.” However, distinct domain-specific patterns were observed: archaeal communities were dominated by a small number of high-abundance ASVs, resulting in a steep decline and a concentrated tail in the distribution curve; in contrast, bacterial communities showed a more gradual decline and an extended long tail. Rare ASVs (defined as those with relative abundances <0.01%) accounted for 88.15% of the bacterial community, compared to 73.82% in the archaeal community, indicating a more prominent “rare biosphere” in bacteria and a closer alignment with stochastic assembly. Archaeal communities, on the other hand, appeared to be more strongly shaped by selective pressures, exhibiting greater deviation from neutrality.

**Fig. 3.**
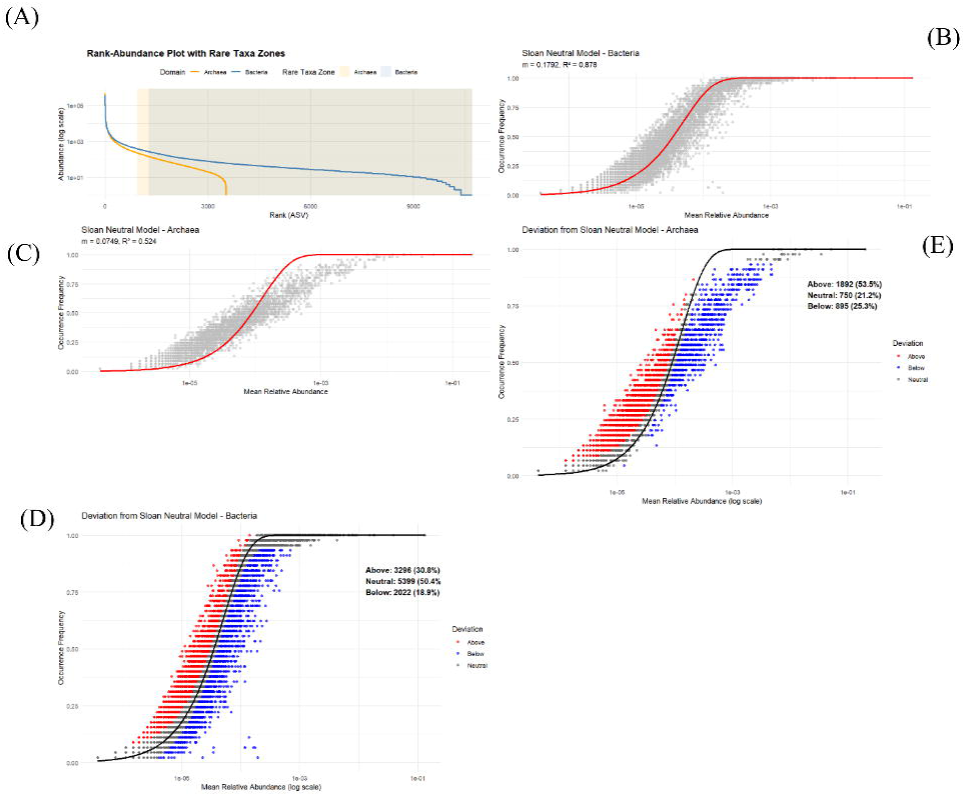
Neutral model fitting and ASV-level distribution across bacterial and archaeal communities. (A) Rank-abundance plots of ASVs for both domains. (B–C) Sloan’s neutral model fits for bacterial (B) and archaeal (C) communities. (D–E) Classification of ASVs as above, neutral, or below the neutral expectation for bacteria (D) and archaea (E).

At the model-based inference level, Sloan’s neutral model was successfully fitted to both datasets. For the bacterial community, the estimated immigration rate (m) was 0.1792, with a model fit R² of 0.878 (Fig. 3B); for archaea, m was 0.0749 with an R² of 0.524 (Fig. 3C). Bootstrap resampling (1,000 iterations) showed narrow 95% confidence intervals for both m and R² estimates, specifically [0.1768, 0.1815] and [0.8687, 0.8855] for bacteria (Fig. S2), and [0.0724, 0.0776] and [0.4859, 0.5531] for archaea (Fig. S3), indicating parameter stability but marked differences in explanatory power.

At the comparative level, we assessed deviations between observed ASV occurrence frequencies and model predictions, classifying ASVs as Above, Neutral, or Below the neutral expectation. Most bacterial ASVs fell within the Neutral category, indicating strong concordance with the model (Fig. 3D), while archaeal ASVs were more widely distributed across the Above and Below categories, suggesting a higher degree of deviation from neutrality (Fig. 3E).

Taken together, both bacterial and archaeal communities showed some degree of neutrality; however, bacterial communities exhibited greater model fit and internal consistency, indicating that stochastic processes like dispersal and drift play a dominant role. In contrast, the weaker neutrality fit and greater deviations in archaeal communities point to stronger deterministic influences, such as environmental filtering. These results support the first DSSDI hypothesis: in dynamic intertidal systems, bacterial assembly is largely stochastic, while archaeal assembly is more deterministic, reflecting domain-specific divergence.

### Spatiotemporal Structuring by Temporal and Environmental Filters

To test the hypothesis 2 that microbial communities in intertidal water environments are structured by spatial heterogeneity, temporal dynamics and environmental variables, we adopted a multivariate framework combining ordination, statistical testing, and variance partitioning. Community composition was first visualized using non-metric multidimensional scaling (NMDS) based on Bray–Curtis dissimilarity. Archaeal communities exhibited clear temporal clustering across the five sampling months (July to November 2022), while spatial separation among the nine sites (A–J) was not evident. In contrast, bacterial communities lacked discernible grouping across both temporal and spatial gradients, suggesting a more diffuse structuring. The ordination quality was adequate for both domains (stress = 0.0067 for Archaea; 0.1139 for Bacteria) (Fig. 4A and Fig. 4B).

**Fig. 4.**
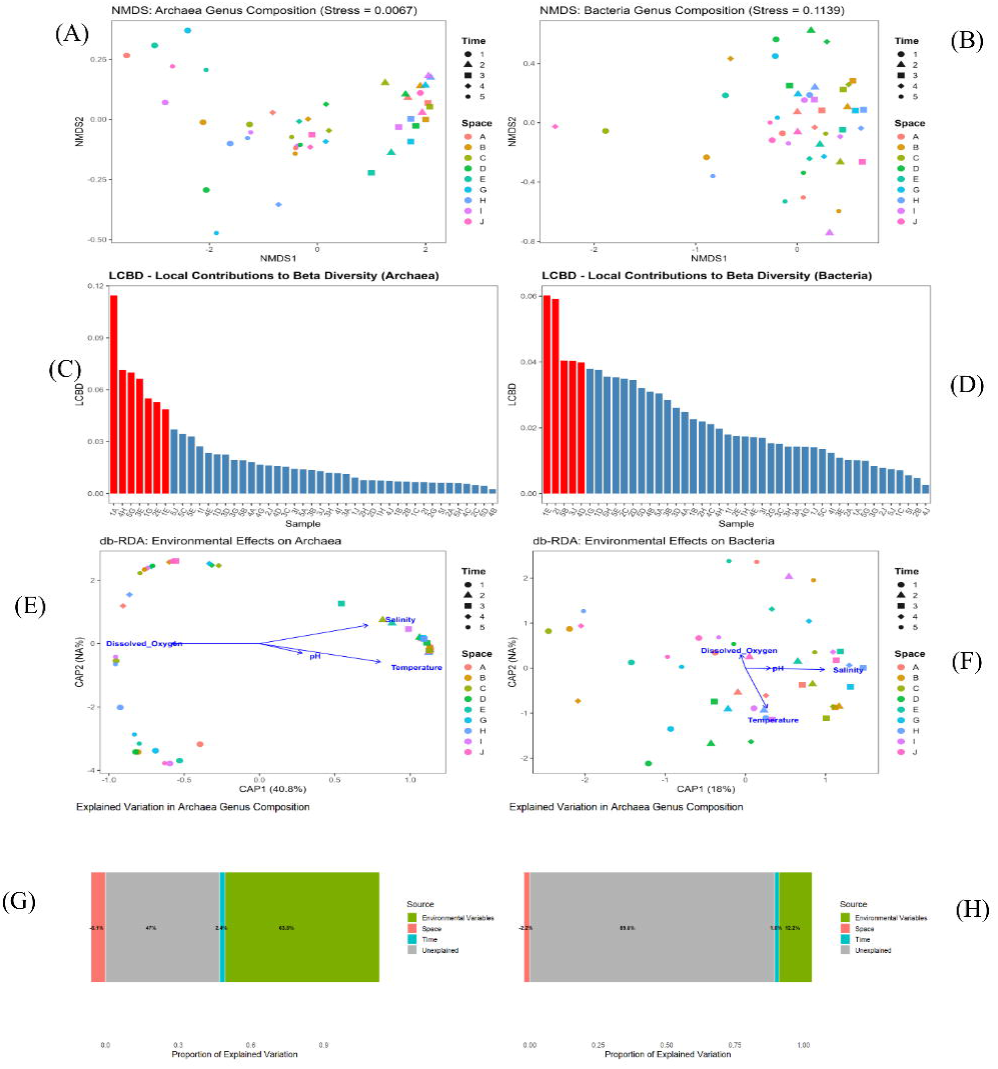
Ordination and variation partitioning of microbial community composition. (A– B) Non-metric multidimensional scaling (NMDS) plots of archaeal (A) and bacterial (B) communities based on Bray–Curtis dissimilarity. (C–D) Local contribution to beta diversity (LCBD) values for archaeal (C) and bacterial (D) samples. (E–F) Distance-based redundancy analysis (db-RDA) for archaeal (E) and bacterial (F) communities using environmental variables. (G–H) Variation partitioning analysis of environmental, temporal, and spatial factors for archaeal (G) and bacterial (H) communities.

To identify compositional uniqueness among samples, local contributions to beta diversity (LCBD) were calculated. Seven archaeal samples—1A, 1E, 1G, 2E, 3E, 4H, and 5G—exhibited significantly elevated LCBD values (p < 0.05), suggesting strong temporal or site-specific shifts in community composition (Fig. 4C). These samples were predominantly concentrated in early (T1) and mid (T3–T4) phases of the sampling period. Bacterial LCBD values were more evenly distributed, with only five significant samples (1E, 2I, 3J, 4D, and 5B), lacking clear temporal or spatial coherence (Fig. 4D).

To formally test the influence of explanatory variables, PERMANOVA was performed using space, time, and four environmental factors (temperature, pH, salinity, and dissolved oxygen). For archaeal communities, time explained 13.9% of the variation (R² = 0.139, p = 0.0012), while space was not significant (R² = 0.054, p = 0.9999). Among environmental variables, only salinity had a significant effect (R² = 0.805, p = 0.0021); temperature, pH, and DO were not significant. For bacterial communities, pH emerged as the only significant factor (R² = 0.3298, p = 0.0443), while both time (R² = 0.0436, p = 0.0601) and space (R² = 0.173, p = 0.5832) were non-significant.

To determine whether the observed differences were influenced by heterogeneity in dispersion, betadisper tests were performed. In archaeal communities, temporal groups differed significantly in their multivariate dispersion (p = 0.0176), whereas spatial groups did not (p = 0.994). For bacterial communities, no significant dispersion differences were observed across either grouping factor. Complementary Mantel tests between Bray–Curtis dissimilarity and geographic distance showed no significant spatial autocorrelation (Archaea: r = –0.0420, p = 0.915; Bacteria: r = –0.0671, p = 0.892), further indicating that spatial proximity did not structure microbial composition in either domain.

To assess the role of environmental filtering, distance-based redundancy analysis (db-RDA) was used. The model was highly significant for Archaea (p < 0.0001) (Fig. 4E), supporting the influence of environmental gradients on community structure. For Bacteria, the db-RDA result was marginally significant (p = 0.0053), indicating weaker environmental control (Fig. 4F). Finally, variation partitioning analysis (VPA) quantified the unique and shared contributions of space, time, and environment. For Archaea, environmental variables uniquely explained 63.6% of the variation, with time contributing 2.4% and space contributing negatively (–6.1%), while residual variation remained at 46.9% (Fig. 4G). In contrast, Bacteria exhibited a dominant unexplained component (89.5%), with minor contributions from environment (12.2%), time (1.6%), and space (–2.2%) (Fig. 4H).

These findings support the second DSSDI hypothesis: archaeal communities are shaped primarily by environmental filtering and temporal variation, reflecting deterministic assembly, while bacterial communities show greater stochasticity and weaker environmental associations. This underscores domain-specific divergence in microbial assembly mechanisms.

### Cross-Domain Positive Interactions and Modular Network Structures Enhance Stability

To test the hypothesis (H3) derived from the DSSDI model—that microhabitat sharing and neutral dispersal processes promote stable, positively correlated cross-domain modules—we constructed a consensus co-occurrence network based on genus-level abundance data of the top 39 archaeal and bacterial taxa. Three inference methods (SparCC, SPIEC-EASI, FlashWeave) were integrated to ensure cross-method consistency and statistical robustness. The resulting network comprised 65 nodes and 170 stable edges, of which 82.4% (n = 140) were positive associations between archaeal and bacterial genera. Notably, no within-domain (archaea–archaea or bacteria–bacteria) links remained after intersection filtering, indicating that reproducible co-occurrence patterns are predominantly cross-domain and synergistic in nature.

Topological analysis further revealed a strongly centralized structure, with a highly skewed degree distribution (coefficient of variation = 1.40). Among the top 10 most connected nodes, 8 were archaeal genera, including Methanofollis, Methanomicrobium, Woesearchaeota Incertae Sedis AR16, and Haloferax, highlighting the dominant structural role of archaea in cross-domain modules. This finding was corroborated by k-core analysis, which showed that the majority of high-degree nodes were concentrated in the network core (k ≥ 3) (Table 1).

Community detection using the Louvain algorithm identified 8 distinct modules (modularity Q = 0.252). Stable modules were primarily composed of archaeal core taxa co-occurring with select bacterial genera such as Aestuariivita and Marinimicrobia genera incertae sedis, forming ecologically cohesive cross-domain subnetworks (Fig. 5A). To assess network robustness and reproducibility, we performed 20 bootstrap resamplings (80% of samples per replicate). A total of 86 edges were recovered in ≥70% of replicates (Fig. 5B), with the top 20 edges showing 100% stability. Six hub taxa— including Methanofollis, Methanomicrobium, Woesearchaeota AR16, Haloferax, Marinimicrobia g.i.s., and Aestuariivita—consistently emerged as top-degree hubs in ≥70% of bootstrap networks (Fig. 5C), underscoring the statistical resilience of the network structure.

**Fig. 5.**
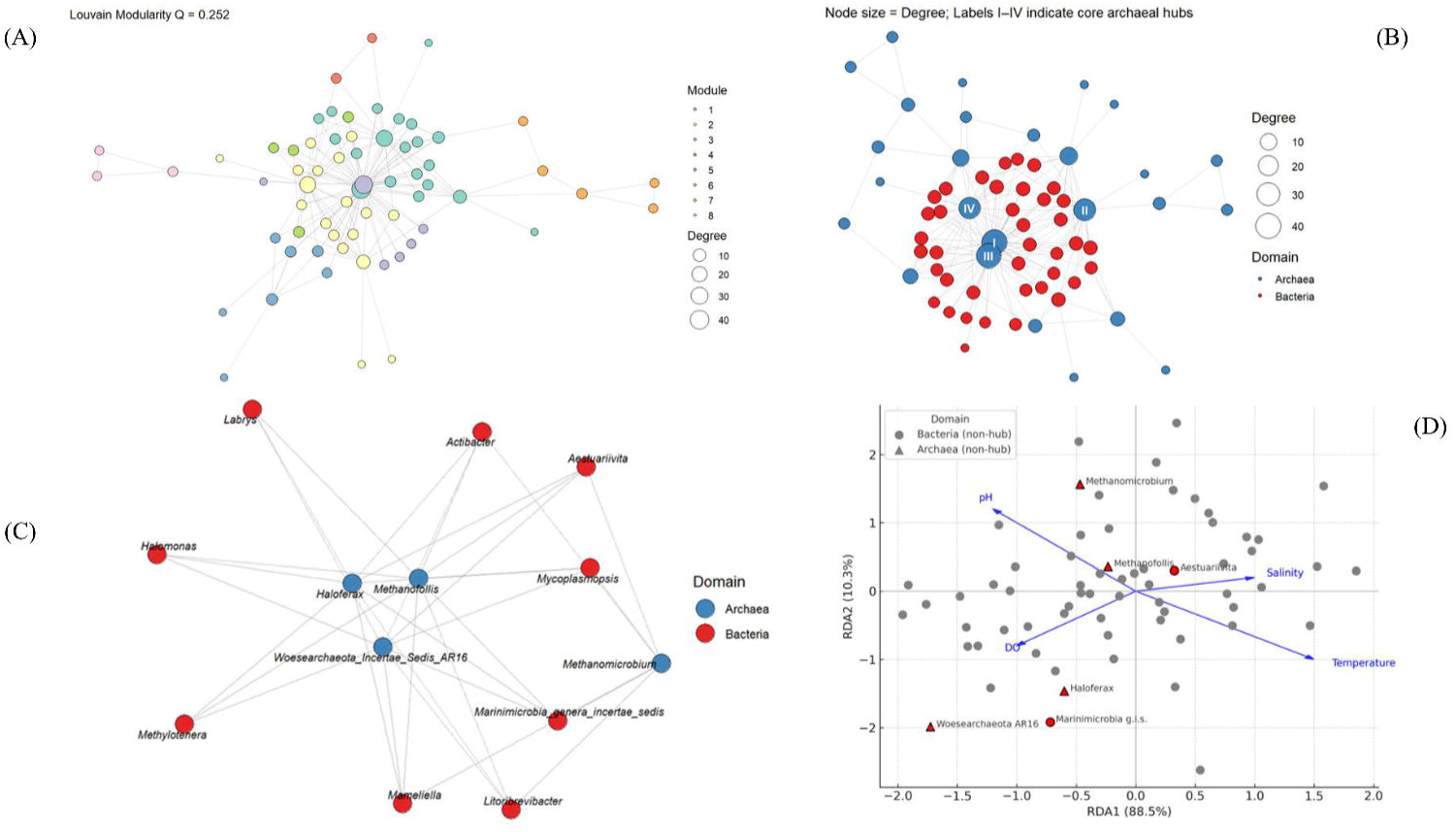
Cross-domain genus-level co-occurrence network and centrality analysis. (A) Consensus network of 39 archaeal and 39 bacterial genera constructed using SparCC, SPIEC-EASI, and FlashWeave; modules identified via Louvain clustering. (B) Edge stability across 20 bootstrap replicates. (C) Degree centrality and hub taxa identification across replicates. (D) Redundancy analysis (RDA) linking hub taxa to environmental variables.

Finally, we performed redundancy analysis (RDA) to evaluate the ecological coherence of the observed modules. The first two canonical axes explained 88.5% and 10.3% of constrained community variation, respectively. Significant environmental drivers included pH, dissolved oxygen (DO), salinity, and temperature—all aligning with the distributional gradients of the six hub taxa (Fig. 5D). Archaeal hubs such as Methanofollis and Methanomicrobium were associated with temperature and salinity, whereas bacterial hubs like Marinimicrobia g.i.s. and Aestuariivita aligned with pH and DO. These results suggest that the core cross-domain modules are not only topologically and statistically stable, but also exhibit shared environmental niche preferences.

These findings support the third DSSDI hypothesis: positive, cross-domain interactions drive modular community architecture that is environmentally structured and statistically robust, consistent with the prediction that such interactions enhance network stability under fluctuating conditions.

### Cross-Domain Functional Complementarity Enhances Ecosystem Functional Resilience

To assess whether functional resilience is maintained despite taxonomic turnover, we predicted metabolic functional profiles based on ASV tables using PICRUSt2. A total of 1,230 functional pathways were inferred for bacterial communities and 980 for archaeal communities.

Principal component analysis (PCA) (Fig. 6A) of KEGG Orthology (KO)-level profiles revealed clear domain-specific functional differentiation. Bacterial functional profiles were more widely dispersed, indicating broader functional diversity, while archaeal profiles were tightly clustered, reflecting narrower metabolic specialization. A Venn diagram of predicted MetaCyc pathways further highlighted functional partitioning: 245 pathways were unique to bacteria, 10 to archaea, and only 119 shared across domains (Fig. 6B). This limited overlap (∼24%) suggests strong cross-domain task differentiation, where bacterial and archaeal communities contribute complementary functions.

**Fig. 6.**
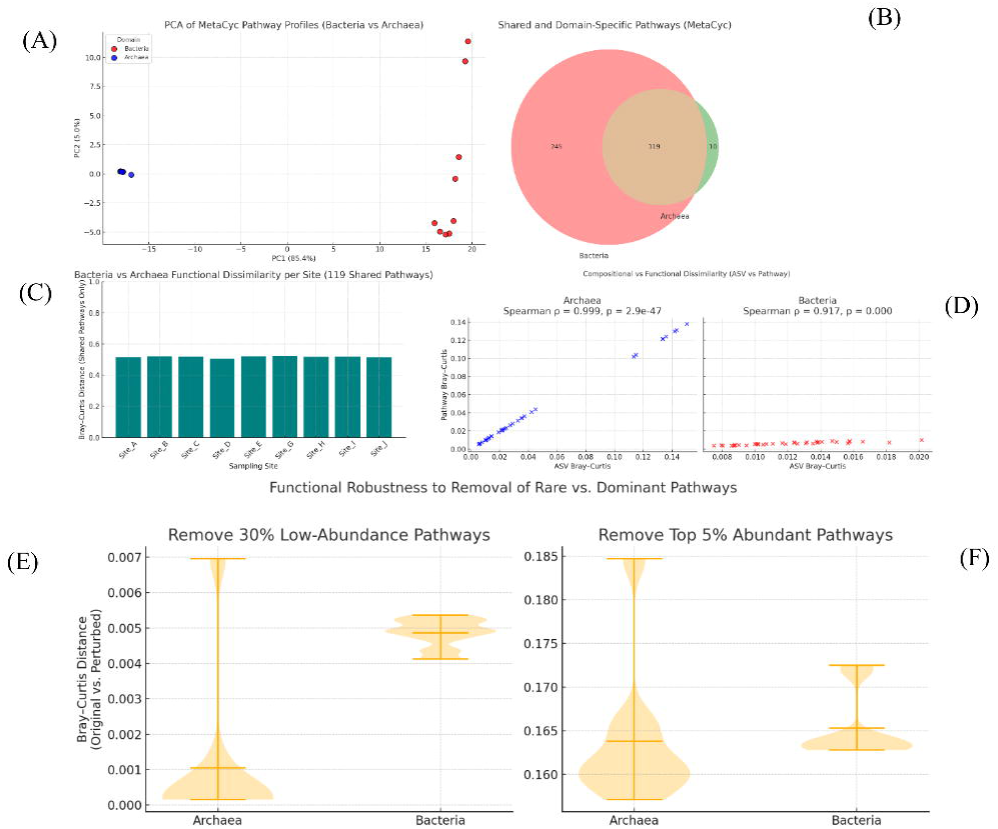
Functional profile comparisons and robustness simulations for bacterial and archaeal communities. (A) Principal component analysis (PCA) of predicted KEGG Orthology-based functional profiles. (B) Venn diagram of shared and unique MetaCyc pathways between bacterial and archaeal communities. (C) Bray–Curtis dissimilarities of shared functional pathways across sampling sites. (D) Correlation between taxonomic and functional Bray–Curtis dissimilarities. (E) Functional profile variation after random removal of rare ASVs (lowest 30% by abundance). (F) Functional profile variation after removal of the top 5% most abundant pathways per sample.

Despite sharing these 119 pathways, Bray–Curtis dissimilarities of their relative abundances remained consistently high across all sampling sites (range: 0.51–0.52) (Fig. 6C), indicating persistent differences in pathway-level contributions between domains. This pattern supports the idea of functional complementarity—where cooperation between bacteria and archaea arises through differentiated but mutually reinforcing roles.

Correlations between ASV-level and pathway-level Bray–Curtis dissimilarities revealed domain-specific patterns of redundancy. In archaea, compositional and functional dissimilarities were nearly perfectly correlated (Spearman ρ = 0.999), suggesting low redundancy and strong dependency of function on specific taxa. In contrast, bacterial communities showed a weaker but still significant correlation (ρ = 0.917), with functional profiles remaining more stable than taxonomic ones (Fig. 6D). This decoupling suggests that bacterial functions are buffered by internal redundancy, allowing ecosystem-level stability under compositional turnover.

To test functional robustness, we performed two disturbance simulations. First, we randomly removed the lowest 30% of ASVs by mean abundance and re-inferred pathway profiles across 100 iterations. Functional profiles remained virtually unchanged (Bray–Curtis: bacteria = 4.31 × 10⁻¹⁷, archaea = 3.20 × 10⁻¹⁷; SD < 5 × 10⁻¹⁷) (Fig. 6E), indicating that rare taxa have negligible impact on core functional capacities. Second, we removed the top 5% most abundant pathways per sample. Even after targeting dominant functions, dissimilarities remained low (bacteria = 0.0038; archaea = 0.0026) (Fig. 6F), confirming that functional architecture is resilient to both rare and dominant component loss.

Functional co-occurrence network analysis revealed four major cross-domain modules via Louvain community detection, each integrating bacterial and archaeal pathways (Fig. 7). Module 1 included 151 bacterial and 38 archaeal pathways, mainly related to glycolysis and amino acid metabolism (Fig. 7A). Module 2 (130 bacterial, 13 archaeal) was enriched in fatty acid biosynthesis and stress response (Fig. 7B). Module 3 (75 bacterial, 48 archaeal) was centered on nucleotide and cofactor biosynthesis (Fig. 7C), and Module 4 (30 archaeal, 8 bacterial) focused on aromatic compound degradation and alternative carbon metabolism (Fig. 7D). These modules suggest a metabolically integrated division of labor across domains, tailored to support biogeochemical resilience under environmental fluctuation.

**Fig. 7.**
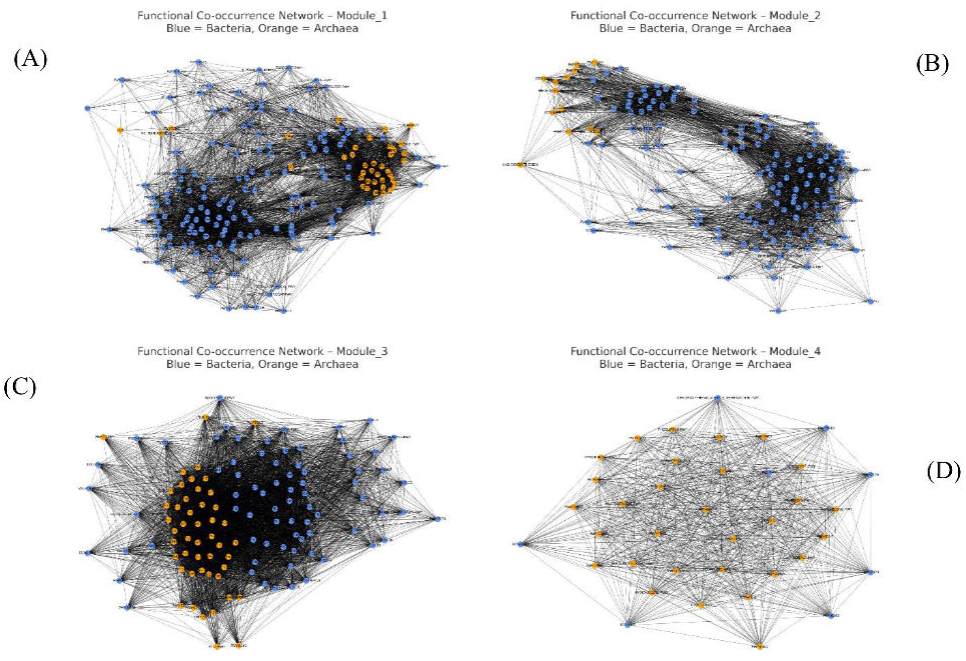
Cross-domain functional co-occurrence network modules. (A–D) Louvain clustering-based functional modules inferred from pathway co-occurrence networks. Each panel represents a distinct module integrating bacterial and archaeal pathways: (A) Module 1 is enriched in glycolysis and amino acid metabolism; (B) Module 2 in fatty acid biosynthesis and stress response; (C) Module 3 in nucleotide and cofactor biosynthesis; (D) Module 4 in degradation of aromatic compounds and alternative carbon metabolism.

Taken together, these findings provide strong support for Hypothesis 4 (H4) of the DSSDI framework. Despite significant taxonomic turnover, microbial ecosystems maintain functionally differentiated and cooperative architectures across domains. This domain-integrated modularity enhances system robustness and underscores the predictive utility of DSSDI for understanding functional resilience in complex, dynamic environments.

## Discussion

### Domain-Specific and Context-Dependent Assembly Mechanisms (H1)

Traditionally, microbial ecology has often framed community assembly as a dichotomy driven either by neutral or deterministic forces. However, our study introduces a more nuanced perspective through the DSSDI model—highlighting that different microbial domains respond asymmetrically to these forces. This asymmetry is not merely a matter of methodology or spatial scale but reflects fundamental differences between bacteria and archaea in terms of metabolic breadth, dispersal potential, and environmental adaptability [7, 44–45].

Our findings reveal that in dynamic intertidal systems, bacterial community composition more closely aligns with predictions from neutral theory, suggesting high immigration rates and strong adaptability to environmental fluctuations. This is consistent with the broader "functional plasticity space" observed in bacteria and is also evident in other marine and estuarine systems [9,13,46].

In contrast, archaeal communities exhibit strong environmental filtering signals, particularly influenced by salinity. This aligns with their highly specialized metabolic traits, such as methanogenesis and ammonia oxidation. Prior research has shown that such metabolic specialization can severely constrain ecological breadth, reducing archaeal adaptability under fluctuating conditions [44,47]. Recent phylogenomic studies further suggest that many archaeal clades originated through specific gene acquisitions from bacteria, leading to tighter niche associations and restricted plasticity [48,49].

Modeling bacteria and archaea jointly may thus obscure these fundamental physiological and evolutionary asymmetries. This distinction, importantly, is not static but context-dependent. In regions with low salinity variability, archaeal community composition remains relatively stable. However, under steep salinity gradients, archaeal sensitivity increases markedly—indicating a low tolerance for abrupt environmental changes [45].

Together, these observations support a theoretical advancement: microbial community assembly should be conceptualized as a domain-specific and scale-dependent dynamic process. Conventional universalist assumptions—such as the idea that all microbes follow neutral theory—must be revisited.

These findings suggest that the widespread assumption of taxonomic uniformity in microbial assembly models may obscure key ecological and evolutionary dynamics. DSSDI reframes community assembly not as a single-axis continuum, but as a domain-partitioned process governed by both phylogenetic legacy and environmental context. This conceptual repositioning extends beyond intertidal ecosystems and may hold relevance for other systems where microbial taxa diverge significantly in metabolic capacity, ecological plasticity, or niche specialization—such as freshwater lakes, peatlands, or host-associated microbiomes. We propose that domain-specific assembly asymmetry is not an exception but a generalizable principle shaping microbial community dynamics across ecologically heterogeneous environments.

### Microbial Communities Structured by the Interaction of Spatial Heterogeneity, Temporal Dynamics, and Environmental Gradients (H2)

Environmental heterogeneity is a cornerstone concept in ecological theory, long recognized as a primary driver of biodiversity and community differentiation [50]. Yet in microbial systems, this principle has largely been explored at macroecological scales—such as biogeographic zones—while micro-scale heterogeneity and its interaction with domain-specific traits have received comparatively less attention. Our findings expand this framework by providing empirical support for the DSSDI model’s second hypothesis (H2), which posits that fine-scale environmental variation plays a central role in microbial assembly, particularly under dynamic boundary conditions.

Despite geographic proximity, intertidal bacterial communities exhibited marked divergence and high β-diversity. These differences persisted even in the absence of overt macro-environmental gradients, indicating that localized factors—such as pH microsites, salt crusts, or nutrient hotspots—may serve as dominant filters [51,52]. This observation aligns with prior studies in soils, aquifers, and wetlands, which show that micro-scale spatial niches can outweigh large-scale abiotic gradients in shaping microbial communities [53–55].

In contrast, archaeal communities were temporally consistent and spatially constrained, suggesting a stronger dependence on stable microhabitats. This pattern reflects their narrower ecological amplitude and greater metabolic specialization [44], consistent with the generalist–specialist framework [47]. These differential responses reinforce the DSSDI assumption that assembly is not only domain-specific but also scale-dependent.

Our variation partitioning analyses further support this distinction: environmental variables explained over 60% of archaeal variation but less than 12% of bacterial variation. This implies that bacterial assembly is influenced by unmeasured fine-scale drivers or higher stochasticity, while archaeal assembly is more tightly governed by environmental filtering. Such hidden factors could include microtopography, nutrient microgradients, or biotic interactions [51,56]. Moreover, spatial autocorrelation tests (Mantel tests) revealed no significant distance-decay patterns, challenging the classical assumption that geographic proximity predicts community similarity—especially in highly dynamic systems like coastal zones [53].

Collectively, these results support a shift from macro-scale, gradient-centric paradigms toward a multi-scalar, context-aware understanding of microbial community assembly.

The DSSDI framework formalizes this transition by identifying localized environmental structure as a distinct and influential axis of differentiation. We propose that in fluctuating environments, local microhabitats—not merely broad abiotic conditions—act as dynamic ecological filters, whose effects are mediated through domain-specific traits and strategies. This context-dependent view provides a unifying theoretical lens for interpreting diversity and community structure across both spatial and temporal dimensions.

Finally, the high explanatory power of salinity for archaeal community structure (R² = 0.805) likely reflects the ecological dominance and niche sensitivity of a few key taxa, particularly Nitrosopumilus. Ammonia-oxidizing archaea are known to exhibit strong salinity preferences and narrow environmental tolerances. Therefore, this filtering signal is not an artifact of low richness, but rather a reflection of functional specialization within dominant archaeal lineages.

### Cross-Domain Interactions Promote Modular Structure and Support System Stability (H3)

Ecological network theory highlights modularity as a key structural property that enhances community stability, particularly under disturbance [11,12]. While this concept has been extended to microbial systems, most studies have focused on within-domain co-occurrence, often overlooking the potential of cross-domain interactions. Here, we provide empirical support for the DSSDI model’s third hypothesis (H3): that cooperative cross-domain interactions enhance modular network architecture and confer functional stability.

Our reconstructed genus-level co-occurrence networks revealed that 82.4% of stable edges were positive bacterial–archaeal interactions. Remarkably, filtering out weak links eliminated nearly all within-domain associations, underscoring the pivotal role of cross-domain links in maintaining network cohesion. This pattern supports prior evidence that microbial keystone taxa frequently lie at inter-domain junctions [57,58], where they mediate critical ecological interactions.

Within modules, archaeal taxa such as Methanofollis and Haloferax consistently emerged as core nodes with high connectivity and central k-core scores. These taxa are known to rely on hydrogen, acetate, or reduced nitrogen compounds provided by partner bacteria [59,60], suggesting that the observed network topology reflects not only statistical associations but underlying physiological interdependence.

Importantly, modularity in these cross-domain networks does more than maintain structural integrity—it enhances resistance to perturbation. Theoretical models in ecology suggest that modular networks localize disturbances and limit systemic propagation of shocks [61,62]. Our results align with this principle, indicating that positive bacterial–archaeal modules function as ecological shock absorbers in dynamic intertidal environments.

These findings suggest that cross-domain modularity is not merely an emergent byproduct of microbial interactions but a foundational organizing principle that underpins ecosystem resilience. DSSDI advances ecological network theory by embedding interdomain cooperation within a taxonomic-contextual framework, wherein domain asymmetry and syntrophic linkages jointly stabilize microbial community architecture. This mechanism is likely applicable to other ecosystems where modularity and metabolic complementarity play central roles—such as plant rhizospheres, animal gut microbiomes, and contaminated soils.

To ensure the robustness of our network inferences, we employed a conservative edge-filtering strategy, retaining only those associations supported by all three methods: Spearman correlation, SparCC, and MIC. While this increased confidence in edge stability, it also inherently filtered out weaker within-domain connections. The resulting dominance of cross-domain edges reflects consistently strong interdomain correlations, rather than artefacts of methodological bias. Notably, networks constructed under relaxed thresholds (e.g., ≥2 methods agreement) still showed bacterial–archaeal edges as topologically central (Supplementary Fig. S3), further supporting their structural and functional significance.

### Cross-Domain Functional Complementarity Buffers Compositional Variability and Sustains Ecosystem Functional Stability (H4)

Traditional models of microbial functional stability often rely on the assumption of within-domain redundancy, suggesting that taxonomically distinct species within the same domain can perform interchangeable metabolic roles to ensure functional persistence [17]. However, this assumption has increasingly come under scrutiny, particularly under conditions of environmental stress, where functional plasticity may be insufficient to buffer taxonomic turnover [18]. Our findings offer robust empirical support for the fourth hypothesis of the DSSDI framework (H4): that cross-domain functional complementarity, rather than within-domain redundancy, underpins microbial ecosystem stability in fluctuating environments.

Despite substantial taxonomic variation across sampling points (Bray–Curtis dissimilarity: Bacteria = 0.408; Archaea = 0.310), the predicted functional compositions remained strikingly stable (Bacteria = 0.012; Archaea = 0.045). Principal component analysis further revealed clear functional partitioning between domains: bacterial communities were broadly associated with heterotrophic functions such as organic matter degradation and amino acid metabolism, whereas archaeal communities were functionally specialized in ammonia oxidation and methane metabolism. These observations support a model of cross-domain task differentiation, in which distinct metabolic roles are strategically allocated across bacteria and archaea rather than duplicated within them [63,64].

To verify the robustness of this functional structure, we performed two complementary in silico disturbance simulations. In the first, we removed the lowest 30% of ASVs by mean abundance per site to mimic stochastic taxonomic turnover. The resulting functional profiles remained virtually unchanged, with Bray–Curtis dissimilarities close to zero (Bacteria = 4.31 × 10⁻¹⁷; Archaea = 3.20 × 10⁻¹⁷), indicating the presence of a resilient modular buffering structure that spans taxonomic domains. To address potential concerns about the influence of rare taxa alone, we also performed a supplementary simulation removing the top 5% most abundant pathways per sample. Functional compositions remained comparably stable (Bacteria = 0.0038; Archaea = 0.0026), reinforcing the interpretation that core metabolic stability is not contingent on either dominant or peripheral components, but instead emerges from the overall cross-domain modular architecture. These findings align with emerging work suggesting that microbial multifunctionality is more effectively preserved through domain-integrated metabolic networks than through redundancy within individual taxonomic groups [65,66].

Beyond its theoretical significance, this cross-domain buffering mechanism carries crucial ecological implications in the context of global change. As climate-induced shifts in temperature, salinity, and redox conditions intensify, functionally integrated bacterial and archaeal networks may serve as ecological "safety nets", maintaining consistent ecosystem-level metabolic throughput despite taxonomic instability. This interpretation echoes Cavicchioli et al.’s call for incorporating microbial processes into global climate models [2] and complements Jansson and Hofmockel’s emphasis on microbial configuration as a central determinant of biogeochemical resilience [3].

Taken together, the DSSDI framework reframes microbial resilience beyond the traditional "diversity equals stability" paradigm. Rather than assuming functional interchangeability among taxa, DSSDI posits that modular task allocation and cross-domain interdependence provide the structural basis for maintaining functional stability in dynamic ecosystems. Our findings establish cross-domain functional complementarity not merely as a descriptive pattern, but as a predictive and generalizable mechanism that buffers microbial ecosystems against both stochastic and deterministic perturbations.

### Methodological Considerations

This study primarily relied on 16S rRNA gene-based taxonomic profiling and functional inference. While this approach is widely adopted and informative, it has intrinsic limitations. First, the taxonomic resolution of archaeal lineages is constrained by the incomplete coverage of archaeal genomes in current reference databases, which may lead to underrepresentation or misclassification. Second, functional predictions derived from 16S data—such as those generated using PICRUSt2—are indirect extrapolations based on known genome annotations, rather than direct measurements of gene expression or metabolic activity. As such, they reflect putative functional potential rather than realized function, and may not fully capture dynamic responses to environmental conditions.

To overcome these limitations and move toward mechanistic validation, future studies should incorporate multi-omics approaches such as shotgun metagenomics, metatranscriptomics, and metabolomics. These methods will enhance the resolution and accuracy of microbial functional profiling, and provide direct evidence of metabolic activity and ecological function. Such datasets are essential for evaluating the structural and functional robustness predicted by the DSSDI framework, particularly under fluctuating environmental regimes.

Additionally, while the co-occurrence network analysis used here provides valuable insights into species associations and modular structure, this approach is inherently correlative and cannot infer causal relationships. To rigorously test the mechanistic role of cross-domain interactions in promoting ecosystem stability, future work must include experimental perturbations, trait-informed synthetic communities, or longitudinal mesocosm trials capable of disentangling interaction effects from confounding variables.

Beyond these technical considerations, further methodological development should aim to operationalize DSSDI as a testable and modular framework for cross-domain ecological experimentation. This includes establishing standardized metrics to quantify (i) domain-specific assembly asymmetries (e.g., normalized stochasticity ratio, trait dispersion), (ii) network modularity and robustness (e.g., node deletion simulations), and (iii) taxonomic–functional decoupling (e.g., complementarity vs. redundancy indices). These components would transform DSSDI into a flexible analytical toolkit for hypothesis testing, system comparison, and predictive modeling across ecosystems.

This study represents the first formal articulation of the DSSDI framework, emphasizing its theoretical structure and explanatory power through correlation-based patterns. Although causality is not claimed here, correlation serves as a logical entry point for causal inference. Future iterations of DSSDI may integrate experimental, trait-based, and multi-omics data to probe underlying mechanisms more directly. This modular and extensible architecture distinguishes DSSDI from prior assembly models and positions it as a forward-looking platform for microbial ecological theory that is domain-explicit, interaction-aware, and predictive by design.

To our knowledge, this is the first conceptual and empirical model that explicitly integrates domain-specific assembly mechanisms with interdomain functional cooperation in a unified ecological framework. By formally presenting and validating the DSSDI model, we establish both the theoretical foundation and the publication priority of this approach. We encourage the scientific community to test, expand, and refine the DSSDI framework across different ecosystems and microbial domains.

## Conclusion and Future Directions

In conclusion, the Domain-Specific Stochastic–Deterministic Integration (DSSDI) framework synthesizes domain-specific community assembly mechanisms, stochastic– deterministic dynamics, and cross-domain functional complementarity into a flexible and scalable theoretical model. It advances microbial ecological theory by resolving previously overlooked asymmetries between microbial domains, and provides a predictive basis for understanding community resilience under environmental fluctuation.

While this study validates the DSSDI framework in dynamic intertidal systems, broader empirical testing is needed to assess its robustness across diverse ecosystems— including terrestrial, aquatic, and deep-sea habitats. The conceptual foundations of DSSDI—namely, domain-partitioned assembly rules and interaction-driven functional buffering—are inherently extensible to other microbial domains beyond Bacteria and Archaea, such as fungi, protists, and viruses. This makes DSSDI a promising scaffold for studying cross-kingdom interactions in microbiomes shaped by complex ecological forces.

Moreover, the core principles of DSSDI are likely applicable to ecosystems characterized by strong spatiotemporal variability and physicochemical gradients, such as soils, wetlands, host-associated microbiomes, and human-impacted environments. In such contexts, local assembly rules and interdomain cooperation may critically determine ecosystem function. DSSDI offers a unifying theoretical scaffold for linking microbial assembly dynamics with functional resilience across ecological scales, providing a much-needed predictive lens for understanding microbial contributions in the face of rapid global change.

Looking forward, we envision DSSDI as a practical framework for guiding future microbial ecology research. Key directions include: (1) testing domain-specific assembly asymmetries in extreme or unstable environments (e.g., hypersaline lakes, arid deserts, hydrothermal vents); (2) incorporating metagenomic, metatranscriptomic, and metabolomic data to enhance resolution of functional complementarity and metabolic coupling; (3) expanding to additional microbial groups (e.g., fungi, viruses) to evaluate cross-kingdom applicability; and (4) integrating manipulative experiments

and ecological simulations to determine whether modularity can reliably predict functional resilience. Special attention should also be given to anthropogenically disturbed systems, where domain-level structure and interdomain networks may shift dramatically under pressure.

Ultimately, DSSDI bridges classical community assembly theory, ecological network analysis, and resilience modeling. As both a conceptual lens and a testable framework, it lays the foundation for a new generation of microbial ecology—one that is domain-explicit, interaction-aware, and predictive by design.

## Acknowledgments

We would like to thank Jui-Chun Hsu, Hsuan-Chung Hsieh, Yung-Teng Chang for this work.

## Author Contributions

MT Wan conceived the study and supervised the work. PJ Liu performed the experiments and provided a part of conceptual advice. MT Wan and PJ Liu analyzed the data, and MT Wan wrote the manuscript. All authors reviewed and approved the manuscript.

## Conflict of Interest

The authors declare that this study was conducted in the absence of any commercial or financial relationships that could be construed as potential conflicts of interest.

## Funding

This research project was supported by the HMBio-WANYU Biotechnology Co., Ltd., Taiwan (R.O.C.) (No. HF005-1230-01-WYM).

## Data Availability

Available upon request

## Notes

### Competing Interest Statement

The authors have declared no competing interest.

